# Peroxisomes contribute to intracellular calcium dynamics

**DOI:** 10.1101/2020.09.02.279174

**Authors:** Yelena Sargsyan, Uta Bickmeyer, Katrin Streckfuss-Bömeke, Ivan Bogeski, Sven Thoms

## Abstract

Peroxisomes communicate with other cellular compartments by transfer of various metabolites. However, whether peroxisomes are sites for calcium handling and exchange has remained contentious. Here we generated sensors for assessment of peroxisomal calcium and applied them for single cell-based calcium imaging in HeLa cells and cardiomyocytes. We found that peroxisomes in HeLa cells take up calcium upon depletion of intracellular calcium stores and upon calcium influx across the plasma membrane. Further, we show that peroxisomes of neonatal rat cardiomyocytes and human induced pluripotent stem cell-derived cardiomyocytes can take up calcium in a controlled manner. Our results indicate that peroxisomal and cytosolic calcium signals are tightly interconnected. Hence, peroxisomes may play an important role in shaping cellular calcium dynamics by serving as buffers or sources of intracellular calcium.

## Introduction

Calcium ions (Ca^2+^) play a decisive role in the regulation of many cellular processes and inter-compartment communication, especially in excitable cells like neurons or cardiomyocytes (CMs) (Clapham, 2007). In CMs, for example, cytosolic Ca^2+^ directly engages in cell contraction. At the same time, mitochondrial Ca^2+^ coordinates ATP production and energy demand in CMs (Williams et al., 2015), highlighting the importance of intracellular organelles in Ca^2+^ redistribution. The main sites of Ca^2+^ entry to the cell and intracellular calcium signal regulation are the plasma membrane (PM) and intracellular calcium stores, in particular those of the endoplasmic reticulum (ER) (Paupe and Prudent, 2018).

Excess of organellar Ca^2+^ can be detrimental for health. Elevated mitochondrial uptake increases mitochondrial reactive oxygen species (ROS) production and is associated with heart falure and ischemic brain injury (Santulli et al., 2015; Starkov et al., 2004). Reversely, mitochondrial ROS decreases if Ca^2+^ uptake to mitochondria is suppressed (Mallilankaraman et al., 2012; Tomar et al., 2016). Understanding of principles and mechanisms of organellar Ca^2+^ handling provides starting point to develop interventions in dysregulated calcium handling.

Peroxisomes are small intracellular organelles with a phospholipid bilayer membrane. In concert with evolutionarily conserved functions in lipid and ROS metabolism, peroxisomes are highly plastic and change in their number, morphology and content upon environmental stimuli (Smith and Aitchison, 2013). Communication of peroxisomes with other cellular compartments through exchange of ROS or lipid metabolites is essential for human health (Castro et al., 2018; Schrader et al., 2020; Wanders et al., 2015). Yet, peroxisomal Ca^2+^ has not been studied in excitable cells before, and there are contradicting data about the Ca^2+^ handling in peroxisomes and its dependence on cytosolic Ca^2+^ (Drago et al., 2008; Lasorsa et al., 2008). It has been suggested that peroxisomes are potential targets of Ca^2+^ signaling pathways that initiate outside of the peroxisome or serve as a cytosolic Ca^2+^ buffer, but peroxisomes may also take up Ca^2+^ due to their own need (Drago et al., 2008; Islinger et al., 2012).

Measurement of Ca^2+^ dynamics *in vivo* inside cellular organelles was driven by the development of Ca^2+^-sensitive fluorescent proteins, also known as genetically encoded Ca^2+^ indicators (GECIs) (Gibhardt et al., 2016; Pozzan and Rudolf, 2009). Ca^2+^ dynamics was analysed in the ER, in mitochondria, the cytosol, and in lysosomes by using GECIs (McCue et al., 2013; Whitaker, 2010). GECIs have a Ca^2+^ binding domain, usually calmodulin (CaM). Ratiometric pericam is a single fluorophore-based GECI with circularly permuted EYFP (cpEYFP) as the fluorophore (Nagai et al., 2001). A special role among GECIs play chameleon-based sensors that use Förster resonance energy transfer (FRET). Here, Ca^2+^ results in a conformational change, that decreases the distance between donor (typically CFP) and acceptor (typically a YFP variant) so that FRET occurs (Gibhardt et al., 2016; Palmer and Tsien, 2006; Pérez Koldenkova and Nagai, 2013). The FRET/donor ratio (FRET ratio) correlates with the Ca^2+^ concentration (Figure 1A).

**Figure 1:**
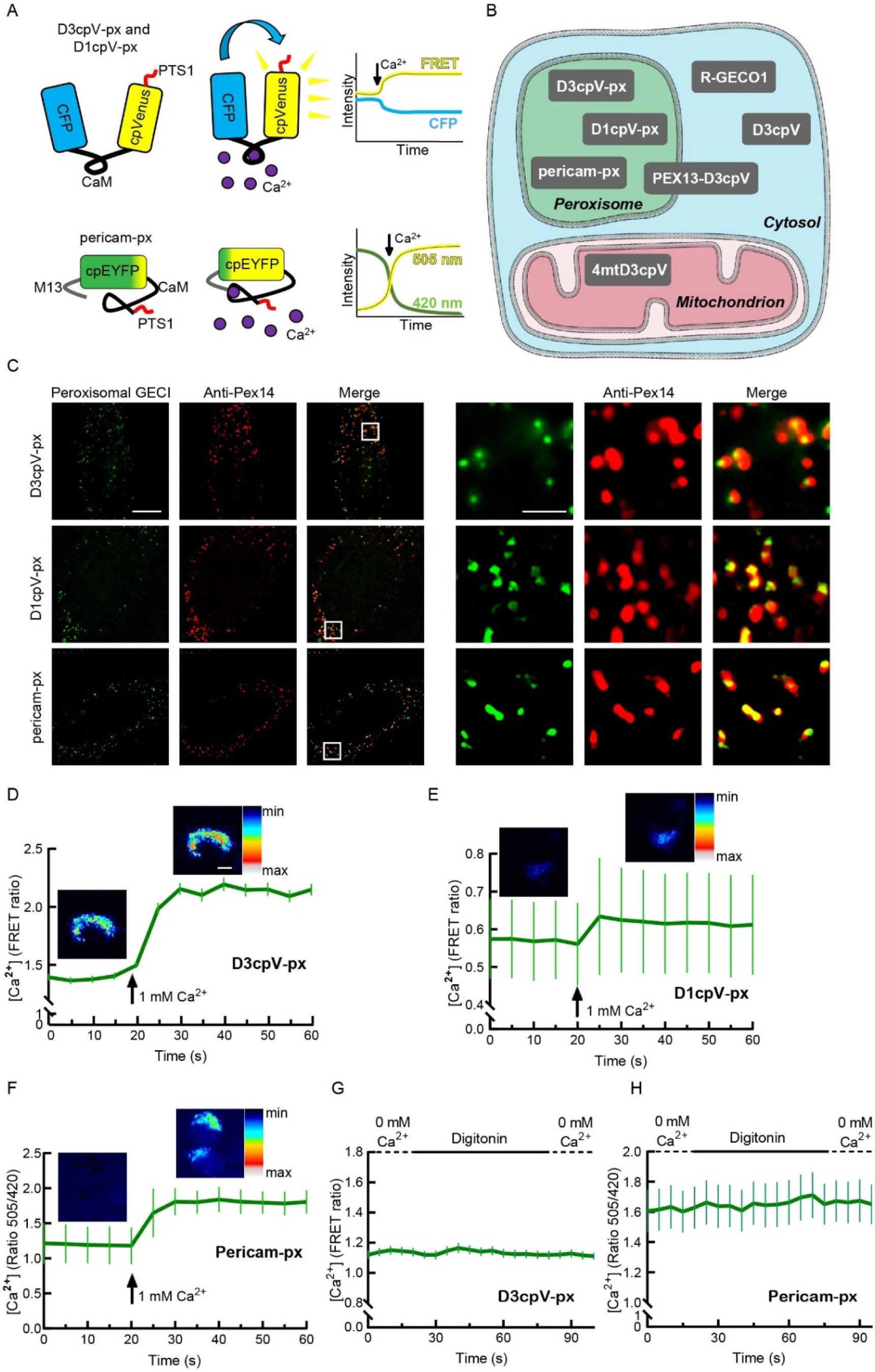
New sensors for peroxisomal Ca^2+^ (A) Genetically encoded calcium indicators (GECIs) targeted to peroxisomes. D3cpv-px and D1cpV-px are FRET sensors with modified CaM sites. Pericam-px is a single-fluorophore based GECI that has M13 and CaM as Ca^2+^ binding sites. In the absence of Ca^2+^, the emission measured when the sensor is excited with 420 nm is higher than when excited with 505 nm. The ratio 505/420 is a measure for the Ca^2+^ concentration. (B) Subcellular localization of GECIs used in this study. (C) Peroxisomal GECIs colocalize with the peroxisomal membrane marker PEX14. HeLa cells were transfected with the GECIs and stained with anti-PEX14 antibodies. The images in the left part of the panel show one cell each (scale bar 10 µm). The cropped areas are marked and magnified in the right part of the panel (scale bar 2 µm). (D-F) D3cpv-px, D1cpv-px, and pericam-px are Ca^2+^ sensitive. Images false-colored with LUT show representative cells before (left) and after (right) Ca^2+^ addition. Curves presented as mean ± SEM. Scale bar: 10 µm. (D) Addition of 1 mM Ca^2+^ to D3cpV-px expressing cells results in 1.5-fold FRET ratio increase, n = 60 cells from three independent experiments. (E) FRET ratio increases 1.08 times when 1 mM Ca^2+^ is added to D1cpv-px expressing cells, n = 33 cells from three experiments. (F) Ca^2+^ addition leads to 1.5-fold increase in 505/420 ratio with Pericam-px, n = 75 cells from three experiments. (G) Measurement of D3cpV-px during cytosol whashout. No change in signal is detected. (H) Measurement of pericam-px during cytosol whashout. No difference of signal before and after cytosol washout is detected, n = 43 cells for D3cpV-px in (G) and n = 45 cells for pericam-px in (H).

The possibility to generate human induced pluripotent stem cells (hiPSCs) from somatic cell sources and to direct their differentiation into almost any cell type make it possible to maintain and study human CMs in culture (Yoshida and Yamanaka, 2011).

This work combines the avantages of organelle-trageted GECIs and hiPSCs. We develop several peroxisomal Ca^2+^ sensors, and we measure intraperoxisomal Ca^2+^ after pharmacological stimulation in non-excitable and excitable cells. We show that peroxisomes take up Ca^2+^ upon cytosolic Ca^2+^ increase both following ER Ca^2+^ store depletion and Ca^2+^ entry to the cells through PM. We also demonstrate that peroxisomes take up Ca^2+^ in rat CMs and hiPSC-CMs.

## Results

### Development and validation of Ca^2+^sensors for peroxisomal Ca^2+^

To assess peroxisomal Ca^2+^, we used three GECIs with different affinities to Ca^2+^: D3cpV, D1cpV and pericam (Figure 1A, Table 1). The sensors were chosen to cover a wide range of K_d_ values to identify the most suitable GECI for intra-peroxisomal measurement. We preferred ratiometric sensors that allow measurements in two wavelengths. This enables direct interpretation of the acquired data by calculating the ratio of intensities at each time point. The ratios provide direct information about Ca^2+^ binding and are independent of the sensor concentration itself (Pérez Koldenkova and Nagai, 2013). For the direct comparison of cellular compartments, we used specific sensors for the cytosol (D3cpV, R-GECO1), mitochondria (4mtD3cpV), and peroxisomes (D3cpV-px, D1cpV-px, pericam-px) (Figure 1B).

**Table 1.** Key properties of the GECIs for cytosol and peroxisome

| Cytosolic GECIs |  |  | Peroxisomal GECIs (this study) |  |
| --- | --- | --- | --- | --- |
| Construct Name | $K_d$ (*in vitro*) | Dynamic range, D | Construct Name | Maximal increase upon 1 mM $\text{Ca}^{2+}$ addition |
| D3cpV | 0.6 $\mu\text{M}$ <sup>(1)</sup> | 5.0 <sup>(1)</sup> | D3cpV-px | 1.50 x |
| D1cpV | 60 $\mu\text{M}$ <sup>(2)</sup> | 1.7 <sup>(3)</sup> | D1cpV-px | 1.08 x |
| Ratiometric-pericam | 1.7 $\mu\text{M}$ <sup>(4)</sup> | 10.0 <sup>(4)</sup> | Pericam-px | 1.50 x |

References: <sup>1</sup> Palmer et al. (2006); <sup>2</sup> Palmer et al. (2004); <sup>3</sup> Greotti et al. (2016) <sup>4</sup> Nagai et al. (2001)

D3cpV is a cameleon-type indicator based on FRET. The conformational change associated with the Ca^2+^ binding to CaM leads to an increase in FRET efficiency and FRET ratio (Pérez Koldenkova and Nagai, 2013). D3cpV has a K_d_ value of 0.6 µM and a dynamic range of 5.0 (Palmer et al., 2006). D1cpV, in comparison, is a FRET sensor with a K_d_ value of 60 µM (Palmer et al., 2004). Finally, pericam is a cpEYFP-based GECI with two excitation peaks at ∼420 nm and 505 nm (Nagai et al., 2001). In the presence of Ca^2+^, a conformational change in the pericam structure shifts the excitation profile so that the 505/420 ratio increases and serves as a measure of Ca^2+^ concentration (Figure 1A). Pericam has a K_d_ value of 1.7 µM and dynamic range of 10 We added strong peroxisomal targeting signals of the PTS1 type to D3cpV, D1cpV, and pericam and tested their localization after transfection by co-staining with antibodies directed against the peroxisomal membrane protein PEX14. All constructs targeted to peroxisomes (Figure 1C).

To test in living cells if D3cpV-px senses Ca^2+^ in the peroxisome, we permeabilized cells by digitonin, washed out the cytosol, and added relatively high Ca^2+^ concentrations. Ca^2+^ addition resulted in drastic increase of FRET and a 1.5-fold increase in FRET ratio (Figure 1D). In order to further illustrate the increase of the FRET signal, we false-colored by using a color look-up table (LUT) the images recorded before and after Ca^2+^the addition (Figure 1D).

When we performed the same type of experiment with D1cpV-px, FRET increased as well after Ca^2+^ addition, showing that the D1cpV-px construct is Ca^2+^ sensitive (Figure 1E). However, following the same stimulation protocol, the signal change of D1cpV-px was only 1.08-fold, and thus considerably smaller than with using D3cpV-px. Due to the low signal change we excluded D1cpV-px from the further experiments on peroxisomal Ca^2+^. Using pericam-px, the third peroxisome-targeted sensor in our experiments, high concentration of Ca^2+^ addition after digitonin treatment resulted in 1.5-fold increase similar as for D3cpv-px (Figure 1F). Based on these results we decided to use D3cpv-px and pericam-px to evaluate Ca^2+^ dynamics in peroxisomes in further experiments.

To study possible mislocalisation or residual signal of peroxisomal Ca^2+^ sensors from the cytosol, we again analysed peroxisomal Ca^2+^ signals following digitonin stimulation of intact cells. In the case of mislocalisation of the sensor to cytosol the signal decrease after digitonin stimulation is expected. We first tested this in D3cpV-px (Figure 1G). There was no signal change observed, suggesting that D3cpV-px has no cytosolic mislocalisation. The cytosol washout also did not change the Ca^2+^signal of the pericam-px before and after digitonin wash, suggesting that pericam-px, like D3cpV-px, is exclusively localized to the peroxisome (Figure 1H).

### Measurement of peroxisomal Ca^2+^in non-excitable cells

We first aimed to compare the maximal possible response of cytosol and peroxisomes to Ca^2+^. On that purpose, we used ionomycin as an ionophore. Ionomycin resulted in fast and immediate increase of cytosolic signal (Figure 2A). Peroxisomal signal increased also, yet, gradually. After reaching its maximum it decreased gradually and in 12 minutes almost returmed to its starting values. The cytosolic reached its half maximal value in the same time with most significant decrese observed in the first two minutes after the maximum. This obeservations suggest that there could aslo be differences in Ca^2+^handling also under near-physiological stimulation.

**Figure 2:**
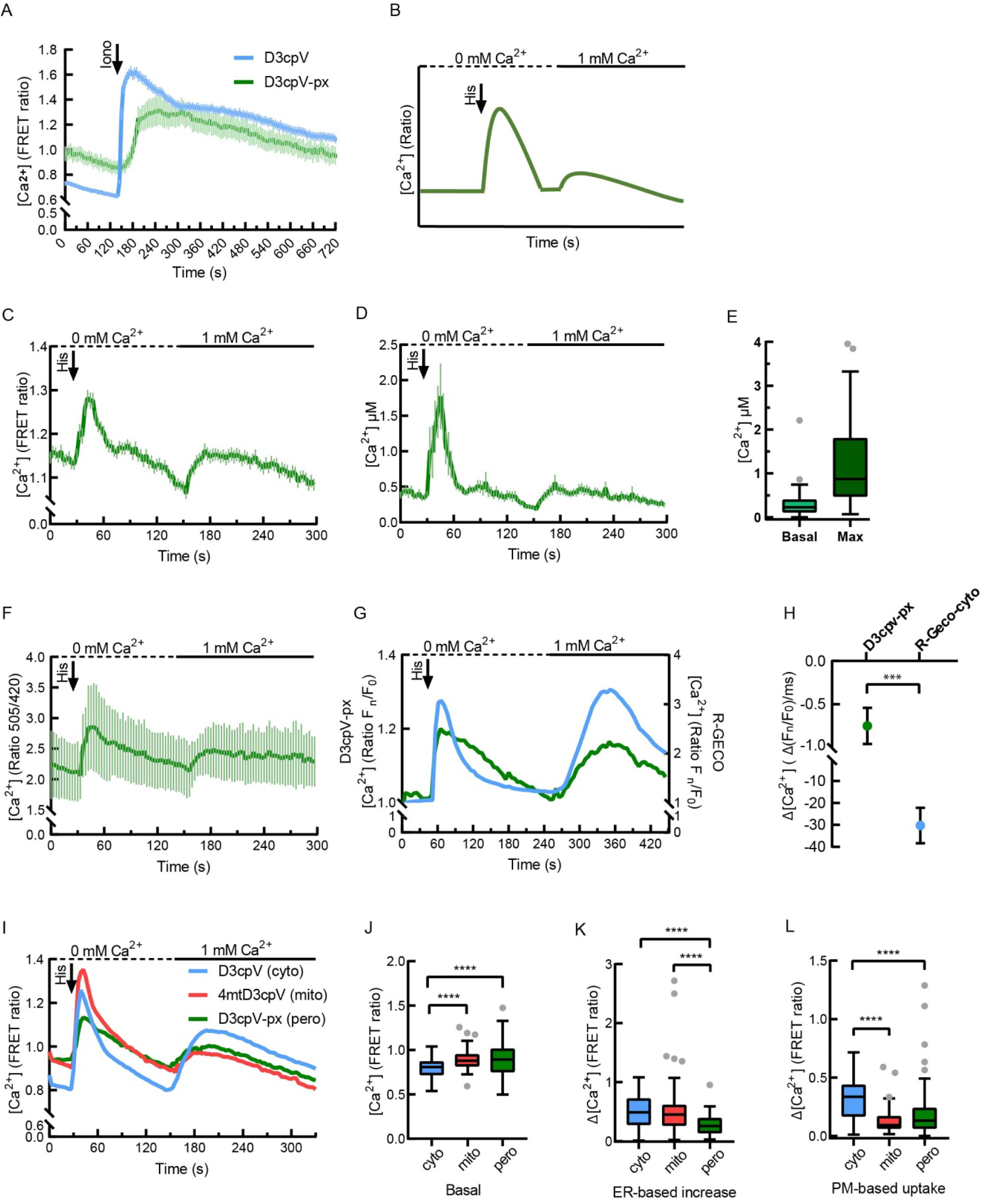
Measurement of peroxisomal Ca^2+^ in HeLa cells. **(A)** Comparison of cytosolic and peroxisomal responces to ionomycin (Iono). In comparison to cytosol, peroxisomal signal increases gradually, n = 16 cells for D3cpV and n = 9 cells for D3cpV-px. (B) Experimental paradigm of a two-step Ca^2+^ measurement in non-excitable cells. 1^st^ peak after histamine (His) addition: ER-store depletion. 2^nd^ peak, after addition of extracellular Ca^2+^: PM-based uptake. (C) Measurement with D3cpV-px according to the paradigm in (B). Two Ca^2+^ peaks of the experimental paradigm are detectable with D3cpV-px, n ≥ 50 cells from three experiments. (D) Absolute Ca^2+^ concentration dynamics calculated from the data in (C). (E) Basal and maximum (max) Ca^2+^ concentrations in peroxisomes based on (C). (F) Measurement with pericam-px according to the paradigm in (B), n = 27 cells from three experiments. (G) Simultaneous measurement of cytosolic (blue) and peroxisomal (green) Ca^2+^. No delay of signal increase after histamine addition, but a delayed drop of the signal in peroxisomes. Left y axis: of D3cpV-px (peroxisomal sensor). Right y axis: F_n_/F_0_ ratio of R-GECO1 (cytosolic sensor), n = 35 cells from three experiments. (H) Decline of F_n_/F_0_ ratio per millisecond (ms) in the linear part of the curves in (G) (from second 65 to 115, Student’s t-test). Kinetic delay in decrease in peroxisomal signal is seen. (I) Comparison of cytosolic, mitochondrial and peroxisomal Ca^2+^ response measured following the paradigm in (B). Characteristic two peaks present in all three compartments. (J) Basal levels of Ca^2+^ in peroxisomes are similar to mitochondria. Analysis performed based on the data from (I). (K) Peroxisomal Ca^2+^ increase upon ER-store depletion is smaller than that of cytosol or mitochondria. Analysis performed based on the data from (I). (L) Peroxisomal Ca^2+^increase upon PM-based cellular uptake of Ca^2+^ is comparable to mitochondria. Analysis performed based on the data from (I). (A, C, D, F) Data presented as mean ± SEM. (J-L) One-way ANOVA followed by Tukey’s post hoc test was used for the statistical analysis. ***p < 0.001, ****p < 0.0001, Tukey’s box plots. Cyto: cytosolic, mito: mitochondrial, pero: peroxisomal. n = 83 (cyto), 116 (mito), 117 (pero) cells from six independent experiments.

Based on the Ca^2+^ measurements in other organelles (Matsuda et al., 2013; Petrungaro et al., 2015; Suzuki et al., 2014; Zhao et al., 2011) we developed an experimental paradigm for peroxisome responses to the depletion and refilling of intracellular Ca^2+^stores, specifically ER, in non-excitable HeLa cells (Figure 2B). The stimulation of cell-surface localized G-protein coupled receptors by 100 µM histamine results in the activation of phospholipase C cascade. Inositol 1,4,5-trisphosphate (IP_3_), the product of the cascade, binds to the IP_3_ receptor on the ER membrane, triggering Ca^2+^ store release. The cells are then exposed to 1 Mm extracellular Ca^2+^, which leads to store-operated Ca^2+^ entry (SOCE) and a second Ca^2+^ elevation in the cytosol. Ca^2+^ is constantly pumped back to the ER through sarcoplasmic/endoplasmic reticulum calcium ATPase (SERCA) (Clapham, 2007).

When we treated HeLa cells expressing D3cpV-px according to this protocol, we observed two peaks (Figure 2C). Histamine addition resulted in a steep and fast increase of intraperoxisomal Ca^2+^ based on depletion of the ER. Addition of extracellular Ca^2+^ resulted in more gradual increase and gradual return to basal levels (Figure 2C).

Using the measurements with D3cpV-px and the known properties of the sensor, we calculated the absolute Ca^2+^ concentration (Figure 2D) applying the formula described by Palmer and Tsien (2006). Under basal conditions, Ca^2+^ level in peroxisomes is around 400 nM and it rises upon near-physiological stimulation with histamine up to 1.8 µM Ca^2+^ (Figure 2E). The Ca^2+^dynamics in peroxisomes measured with D3cpV-px was reproduced by pericam-px: a larger peak is observed after ER-store depletion and a smaller one after extracellular Ca^2+^ addition. The observed ratio curve from pericam-px largely resembles that from D3cpV-px. Since pericam has a K_d_ value of 1.7 µM and covers higher Ca^2+^ concentrations, the observed result confirms the upper limit of peroxisomal Ca^2+^ and the range of Ca^2+^ between 0.4 and 1.8 µM (Figure 2F). However, pericam is described as pH sensitive (Nagai et al., 2001), and since there is currently no consensus regarding pH levels in peroxisomes (Dansen et al., 2000; Jankowski et al., 2001; Waterham et al., 1990) we decided to perform all further experiments with D3cpV-px.

To confirm that the response in our experiments is due to the immediate increase in Ca^2+^ concentration, and to be able to directly compare peroxisomal Ca^2+^ handling with that of the cytosol, cells were co-transfected with D3cpV-px and the mApple-based cytosolic Ca^2+^ sensor R-GECO1, that increases in intensity when binding Ca^2+^ (Zhao et al., 2011). A large increase in the red signal from R-GECO1 was observed both upon ER store depletion and addition of extracellular Ca^2+^ (blue curve in Figure 2G). Although the GECIs used for the measurement in two compartments have different properties that can result in differences in their kinetics, peroxisomes largely follow the Ca^2+^ changes in the cytosol. Interestingly, there is little or no delay between signal increase in cytosol and peroxisomes, and the post-stimulation decline is more gradual and prolonged in peroxisomes compared to the cytosol, indicating the existence of a possible barrier or gate that can be saturated (Figure 2H).

To compare peroxisomal Ca^2+^ levels at rest and under stimulation with that of cytosol and mitochondria, cells were transfected with D3cpV sensors targeting specifically these compartments. FRET ratio was assessed as a direct indicator of Ca^2+^ concentration (Figure 2I-L). All three compartments showed two peaks: one after ER-store depletion with histamine, and another after extracellular Ca^2+^ addition and PM-based uptake (Figure 2I).

The basal levels of calcium in mitochondria and peroxisomes detected with this sensor were comparable and significantly higher than that in the cytosol (typically ≈100 nM, Paupe and Prudent, 2018) in the current settings (Figure 2J). Furthermore, the increase of Ca^2+^ in peroxisomes upon intracellular store depletion with 100 µM histamine was significantly lower than the increase in the cytosol or mitochondria (Figure 2K), speaking against the hypothesis that peroxisomal Ca^2+^ is rising drastically upon stimulation as suggested before (Lasorsa et al., 2008). The addition of extracellular Ca^2+^ resulted in another peak in all three compartments (Figure 2L), evidencing that peroxisomes, like mitochondria depend on the PM-based uptake. Altogether, this suggests that peroxisomes tend to follow Ca^2+^ dynamics of the cytosol.

### Peroxisomal Ca^2+^measurement in cardiomyocytes

We decided to test in neonatal rat cardiomyocytes (NRCMs) the hypothesis that Ca^2+^ can access cardiac peroxisomes. NRCMs are primary cells with a well-developed T-tubule system and serve as a model for electrophysiological studies on CMs (Soeller and Cannell, 1999; Morad and Zhang, 2017).

We adapted the chemical stimulation protocol for the CMs by reducing it to a single stimulation, since the main source of Ca^2+^ in these cells is the ER. We used thapsigargin (Tg) to chemically stimulate the CMs (Figure 3A). Tg is a SERCA antagonist and blocks the constant repumping of Ca^2+^ back to the ER, resulting in Ca^2+^ accumulation in the cytosol. To avoid measurement distortion due to the spontaneous contractile activity of CMs, they were treated with 2,3-butanedione monoxime (BDM) (Gwathmey et al., 1991) before the experiment. As a proof of concept and for direct comparison, we performed the first round of measurements using the cytosol-localized Ca^2+^-sensor D3cpV (Figure 3B-D). A comparison between the Tg-treated cells with the buffer conditions (Figure 3B) demonstrated, as expected, no differences in the basal ratios (Figure 3C), but an increase of cytosolic Ca^2+^ upon Tg addition (Figure 3D).

**Figure 3:**
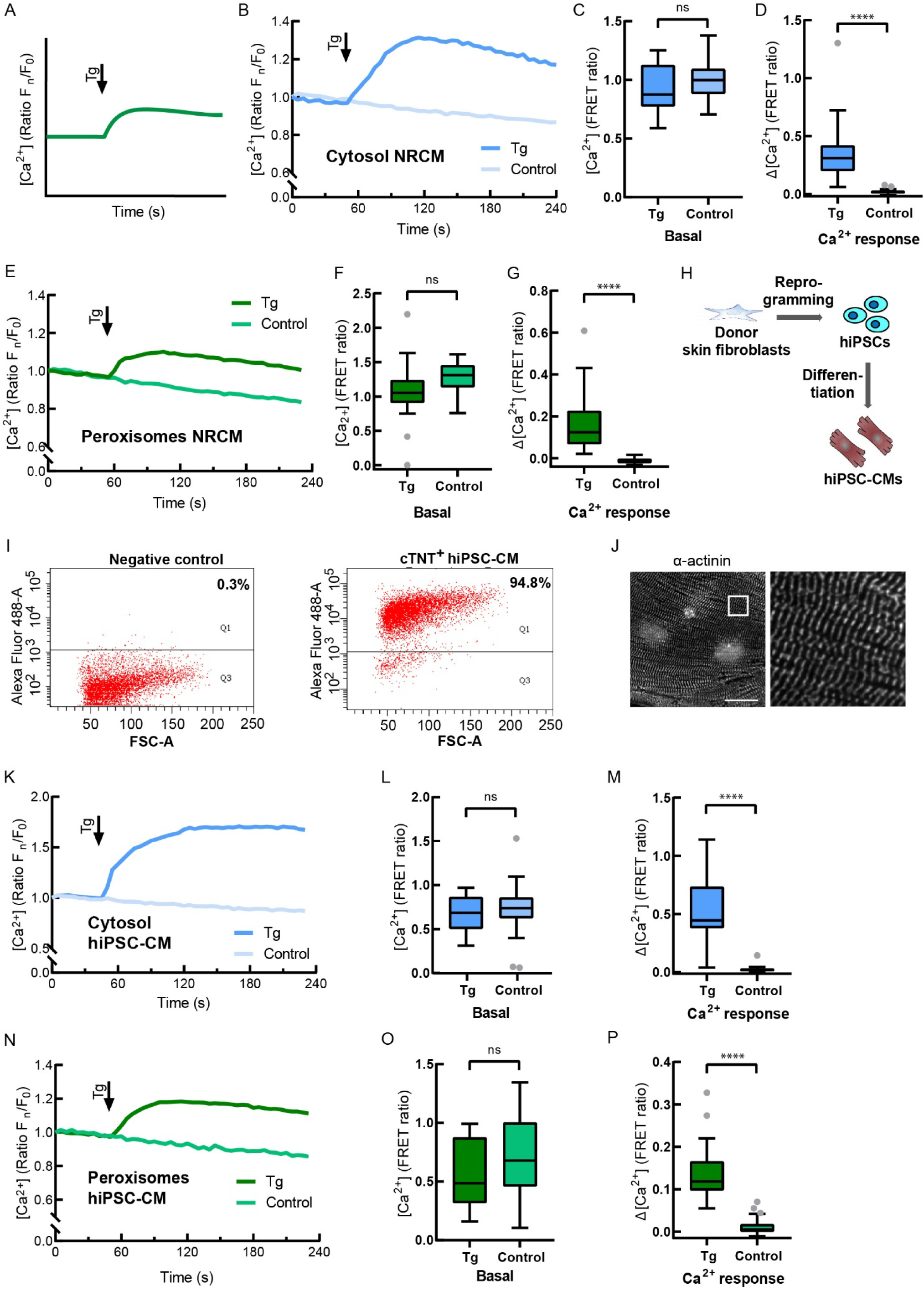
Measurement of peroxisomal Ca^2+^ in cardiomyocytes. **(A)** Experimental paradigm of Ca^2+^ measurement in excitable cells. The peak after thapsigargin (Tg) addition represents Ca^2+^ increase due to the SERCA inhibition and Ca^2+^ retention in the cytosol. (B) Cytosolic Ca^2+^ measurement in NRCMs following the experimental design in (A), n = 25 (Tg), 22 (control) from three experiments. Addition of Tg is compared to the addition of Tg-free buffer (control). (C) Basal levels are not different before the treatment in (B). (D) After Tg addition in (B) cytosolic Ca^2+^ increases. (E) Peroxisomal Ca^2+^ measurement in NRCMs following the experimental design in (A). Addition of Tg is compared to the addition of Tg-free buffer (control), n = 20 (Tg), 31 (control) from three experiments. F) Basal levels of Ca^2+^ are not different before the treatment in (E). (G) Peroxisomal Ca^2+^ increases after Tg addition in (E). (H) HiPSC-CMs generation. Donor skin fibroblasts were reprogrammed to hiPSCs, which were then differentiated to CMs. (I) hiPSC-CMs were stained for cardiac troponin (cTnT) and analyzed by flow cytometry. Negative control without primary antibody. 94.8% of iPSC-CMs are cTnT-positive (cTNT^+^). (J) Immunofluorescence staining visualized α-actinin protein expression and regular sarcomeric organization. Scale bar: 20 μm. (K) Cytosolic Ca^2+^ measurement in hiPSC-CMs with D3cpV following the experimental paradigm for excitable cells in (A). Addition of Tg is compared to the addition of Tg-free buffer (control) to avoid artefacts and false results due to mechanical effect on the cells due to the addition itself. n = 24 (Tg), 27 (control) from three differentiation experiments. (L) No difference is found between two groups before the treatment in (K). (M) Tg addition in (K) results in cytosolic Ca^2+^ increase. (N) Peroxisomal Ca^2+^ measurement in hiPSC-CMs with D3cpV-px following the experimental design in for excitable cells depicted in (A). Addition of Tg is compared to the addition of Tg-free buffer (control). n = 26 (Tg), 33 (control) from three differentiation experimnets. (O) Basal levels of Ca^2+^ are not different before the treatment in (N). (P) Peroxisomal Ca^2+^ increases after Tg addition in (M). (B, E, K, N) Data presented as means from three independent experiments. (C, D, F, G, L, M, O, P) Unpaired Student’s t-test was used for the statistical analysis. ****p < 0.0001, Tukey’s box plots.

To measure peroxisomal Ca^2+^ changes, we transfected NRCMs with D3cpV-px and compared Tg treatment with the untreated control group (Figure 3E). No offset of basal ratios between the two groups was present before treatment (Figure 3F). After the addition of the SERCA inhibitor, peroxisomal Ca^2+^ increased, evidencing peroxisomal Ca^2+^ uptake in NRCMs after store depletion (Figure 3G).

In the next set of experiments, we wanted to know if peroxisomes of human cardiac cells are able to take up Ca^2+^. To test this, human iPSCs created from fibroblasts of a healthy donor were differentiated into CMs using standardized protocols including cardiac mesoderm induction by subsequent activation and inhibition of the WNT pathway (Lian et al., 2013) and metabolic selection (Tohyama et al., 2013) (Figure 3H). Cardiac differentiation was tested for homogeneity by using the cardiac specific marker cardiac troponin T (cTNT) and analysis by flow cytometry at day 90 of differentiation. Our differentiation consisted of 90 %-95 % cTNT-positive cells (Figure 3I). Staining of hiPSC-CMs with antibodies against α-actinin showed a regular sarcomeric striation pattern (Figure 3J).

As a proof of concept and for direct comparison, we measured cytosolic and peroxisomal Ca^2+^ and compared Tg treatment with the addition of Ca^2+^-free buffer without Tg to the control cells (Figure 3K-P). Starting with the same basal ratios as the control samples (Figure 3K and 3L), Tg-treated cells showed a Ca^2+^ increase after the treatment (Figure 3M).

After confirming that Tg can effectively deplete Ca^2+^ stores in hiPSC-CMs, we measured peroxisomal Ca^2+^ in these cells (Figure 3N). No ratio differences were present before Tg treatment (Figure 3O). Ca^2+^-store depletion resulted in an increase of peroxisomal Ca^2+^, confirming peroxisomal Ca^2+^ uptake in hiPSC-CMs (Figure 3P).

To test whether Ca^2+^ enters peroxisomes in a beat-to-beat manner in NRCMs, we field stimulated the cells with 1 Hz frequency (Figure 4A-D). The action potential depolarizes cell membrane resulting in the activation of voltage-gated Ca^2+^ channels in T-tubules (Bootman et al., 2002; Chapman, 1979). As a result, initial minor amount of Ca^2+^ enters the cell. It activates ryanodine receptors on the sarcoplasmic reticulum membrane, resulting into Ca^2+^ release from the stores. Ca^2+^-induced Ca^2+^ release from the stores enables cardiac muscle contraction. During relaxation SERCA and NCX (sodium-calcium exchanger) pump Ca^2+^ back to the intracellular Ca^2+^stores and out of the cells (Clapham, 2007).

**Figure 4:**
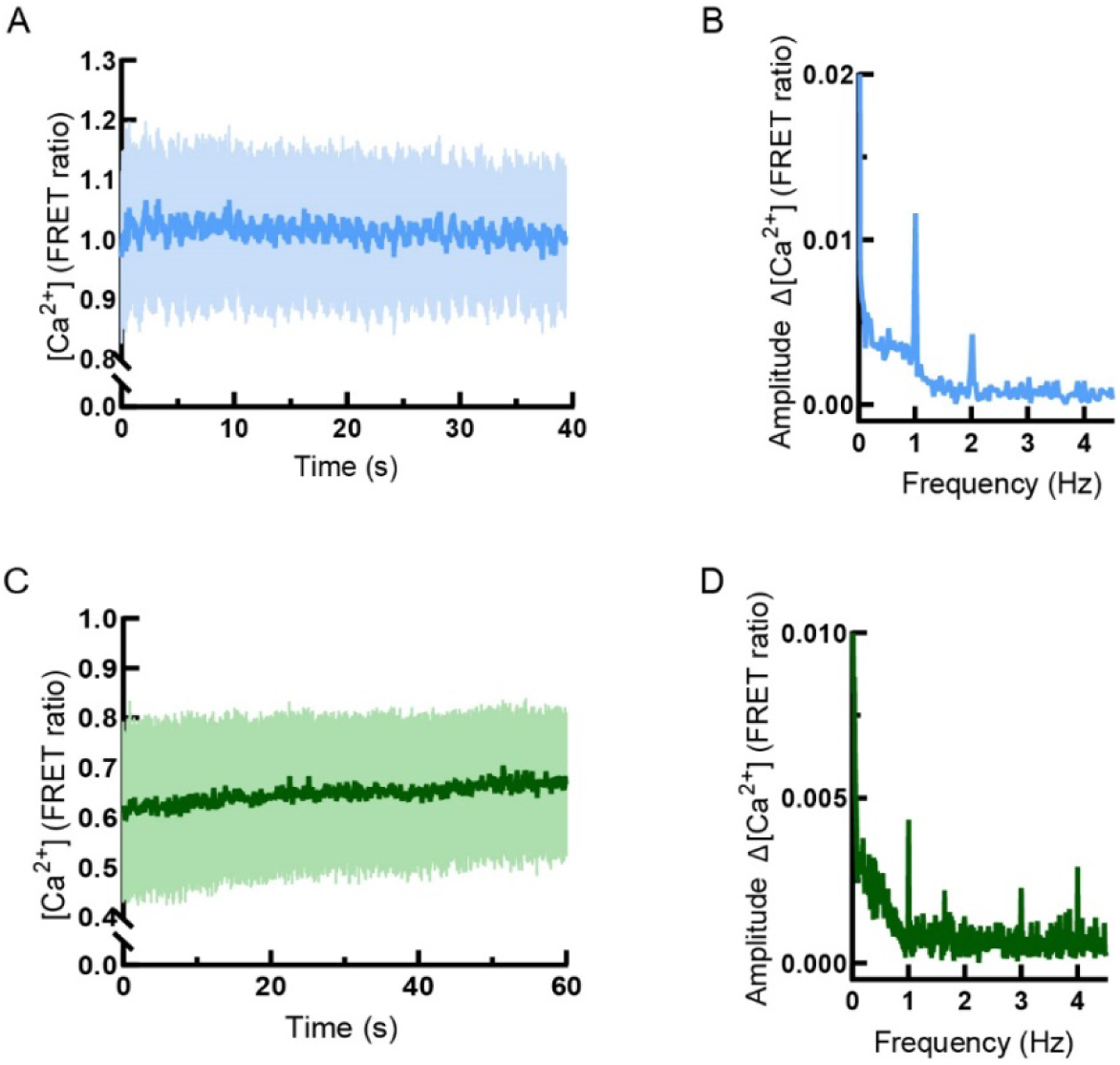
Measurement of peroxisomal Ca^2+^ in paced cardiomyocytes. **(A**) D3cpV transfected NRCMs are stimulated with 1 Hz. Images are taken every 50 ms. Oscillations of FRET ratio are seen, n = 3. (B) FFT from the data in (A). Signal increases are rhythmic and correspond to the pacing frequency. (C) D3cpV-px transfected NRCMs are stimulated with 1 Hz. Images are taken every 100 ms. FRET ratio oscillations are see. n = 3. (D) FFT from the data in (C). Signal increases are rhythmic and correspond to the pacing frequency.

Under field stimulation, we observed rhythmical changes of Ca^2+^ level in the cytosol (Figure 4A). To quantify the amplitude of changes and link to the stimulation we performed fast Fourier transformation (FFT) of the data (Figure 4B). Signal amplitude oscillations in the cytosol were rhythmical and corresponded to the stimulation frequency (Figure 4B).

To test peroxisomal response to electrical stimulation, NRCMs expressing D3cpV-px were paced at a frequency of 1 Hz (Figure 4C). Oscillations observed were smaller in amplitude and appeared less regular than the cytosolic responses. To identify the frequency domain of these oscillations we performed FFT (Figure 4D). The extracted pattern showed amplitude changes at 1 Hz, suggesting that peroxisomes take up Ca^2+^in beat-to-beat manner.

Altogether, these results suggest that peroxisomes in both, rat and human cardiomyocytes were able to take up Ca^2+^ upon intracellular Ca^2+^-store depletion and cytosolic Ca^2+^ increase.

## Discussion

Peroxisomes are metabolically highly active organelles in need of communication with other cellular compartments (Sargsyan and Thoms, 2020). ROS signaling and homeostasis are central to the participation of peroxisomes in signaling pathways (Lismont et al., 2019). In the current work we focused on Ca^2+^ dynamics of peroxisomes as one of the major signaling molecules in the cell. We demonstrate that Ca^2+^ can enter peroxisomes of HeLa cells both when ER-stores are depleted and when cytosolic Ca^2+^ increases after Ca^2+^ entry across the plasma membrane.

Two articles published in 2008 brought forth conflicting data on peroxisomal Ca^2+^. According to Drago et al. (2008), the basal level of Ca^2+^ in peroxisomes equals the cytosolic Ca^2+^ level, whereas Lasorsa et al. (2008) find peroxisomal Ca^2+^ to be 20 times higher than in the cytosol. While Lasorsa et al. (2008) report rise of peroxisomal Ca^2+^ up to 100 µM using an aequorin-based sensor, Drago et al. (2008) suggest slow increase when cytosolic Ca^2+^ rises. Each of the groups used a single yet different technique. These differences in the results can be partially attributed to the different measurement methods and the cell types used. Aequorin imaging requires long incubation times and cell population-based analysis that can be disadvantageous when measuring Ca^2+^ in intracellular organelles. In our experiments with HeLa cells, we found four-fold higher basal peroxisomal Ca^2+^ level compared to the cytosol and increase up to 1.8 µM upon stimulation (Table 1). The range of the changes we report are based on the measurements with D3cpV-px and are supported by the measurement with pericam-px. Hence, we conclude that D3cpV-px can be used for measuring peroxisomal Ca^2+^ concentration in a broad variety of cell types.

Electron microscopic experiments on rodent hearts performed in the 1970s show that peroxisomes are closely associated with T-tubules and with junctional sarcoplasmic reticulum (Hicks and Fahimi, 1977). The sarcoplasmic reticulum is an indispensable site for the excitation-contraction coupling and Ca^2+^ handling in myocytes (Flucher et al., 1994). The localization of peroxisomes to these sites raises the question if cardiac peroxisomes react to Ca^2+^ oscillations on a beat-to beat basis, and/or if they can buffer calcium. HiPSC-CMs provide a wide spectrum of possibilities in cardiac research ranging from drug screening to cardiac regeneration (Yoshida and Yamanaka, 2011). In addition, these cells have been especially used to study patient-specific disease models including arrhythmic disorders and cardiomyopathies demonstrating a robust correlation to the predicted phenotype (Borchert et al., 2017; Liang et al., 2016; Streckfuss-Bömeke et al., 2017).We report here that Ca^2+^ is entering peroxisomes upon intracellular Ca^2+^-store depletion in CMs. Since intracellular store depletion is the main source of Ca^2+^ in CMs in the process of excitation-contraction coupling, it can be hypothesized that peroxisomes take up Ca^2+^ also on beat-to-beat manner in these cells.

Measurement of peroxisomal Ca^2+^ in CMs with FRET sensors in field stimulation suggests that peroxisomal Ca^2+^ increases on beat-to-beat manner. This suggests that peroxisomes may participate in excitation-contraction processes. The exact role of peroxisomes here is the matter of future research. Furthermore, the experimental protocols developed here can be applied to study peroxisomal Ca^2+^ other cell types like neurons.

We found that basal peroxisomal Ca^2+^ levels are higher than cytosolic levels. There are two major ways of generating this Ca^2+^ gradient on the two sides of the membrane. One option could be the energy-dependant uptake mechanism, like SERCA for the ER (Clapham, 2007). We are, however, not aware of any data that can support this model. The second option may be locally high Ca^2+^ concentration at the entry side that would allow more direct channeling of Ca^2+^ (from the ER) into the peroxisomes resulting in relatively high peroxisomal Ca^2+^. This second mechanism is known from the mitochondrial Ca^2+^ handling, where ER-mitochondria contact sites with tethering proteins generate microdomain with locally high Ca^2+^ concentration (Hirabayashi et al., 2017). As a result, Ca^2+^ entry to mitochondria follows the Ca^2+^ gradient but mitochondrial Ca^2+^ is higher compared to cytosol. For the plausibility of the second option for peroxisomes speack the existance of ER-peroxisome contact sites (Costello et al., 2017; Hua et al., 2017). Therefore, we propose a hypothetical model of this mechanism, where most of the components are, however, yet unknown (Figure 5).

**Figure 5:**
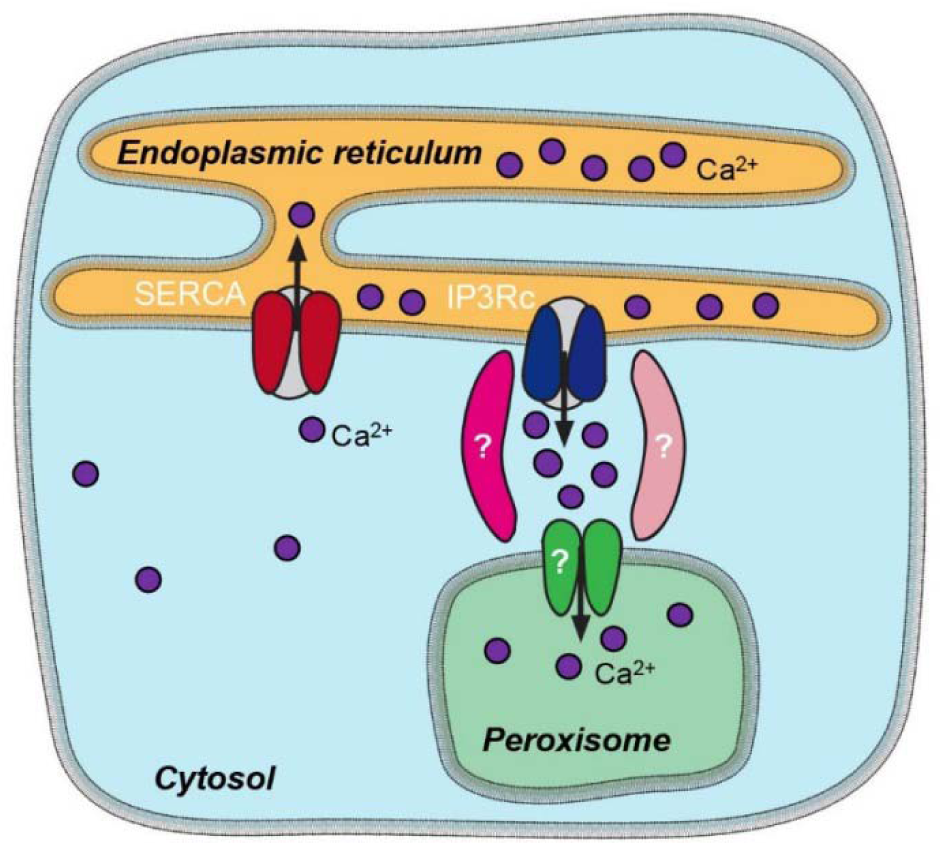
Peroxisomal Ca^2+^ entry and cellular Ca^2+^ distribution. ER Ca^2+^ release triggers Ca^2+^ entry into the peroxisome. In our hypothetical model, ER-peroxisome proximity defines Ca^2+^ microdomains with locally elevated Ca^2+^ concentration shielded from the cytosol. As a result, Ca^2+^ entry to peroxisomes follows the local gradient but peroxisomal Ca^2+^ is eventually higher than in the cytosol. IP3Rc: IP_3_ receptor calcium release channel of the ER.

Although peroxisomal Ca^2+^ levels are higher than cytosolic Ca^2+^ levels, peroxisomes are unlikely to store significant amounts of Ca^2+^ under normal conditions, and they themselves take up Ca^2+^ when intracellular stores are depleted. Under specific conditions, like apoptosis or oxidative stress, the situation may change, however. We show that the rise of peroxisomal Ca^2+^ after histamine stimulation is not delayed and largely follows the cytosolic Ca^2+^. Though there could be a delay due to the binding and conformational changes of GECIs needed before the detection of the increase of the FRET signal, the range of this delay is less than milliseconds and cannot be seen in the experiments described here. We conclude that peroxisomes respond to cytosolic Ca^2+^ since we only found concordant changes of Ca^2+^ concentration in these two compartments.

The question of the cellular function and potential targets of peroxisomal Ca^2+^ is still open. One of the roles of Ca^2+^ could be the regulation of peroxisomal processes. On the other hand, metabolic processes themselves may regulate Ca^2+^ uptake to organelles, as known from mitochondria (Nemani et al., 2020). Whether there is a mutual regulation of metabolic pathways or ROS production localised to peroxisomes is not known. Some plant but not mammalian catalases bind Ca^2+^ (Yang and Poovaiah, 2002). Currently, there are no peroxisomal processes known in mammals that would depend on Ca^2+^. Peroxisomes, however, could serve as an additional cytosolic buffer for Ca^2+^ to take up an excess of cytosolic Ca^2+^ and release it slowly. Based on the findings of this study that the Ca^2+^ concentration in the peroxisome is higher than in the cytosol, it could be that peroxisomes may also serve as additional Ca^2+^ source for the cytosol in extreme situations.

## Methods

### DNA constructs

D3cpV-px (PST 1738) was generated from (pcDNA-)D3cpV (kind gift from A. Palmer and R. Tsien (Palmer et al., 2006) (Addgene #36323)) by amplifying an insert with OST 1599 (GCGCATCGAT GGTGATGGCC AAGTAAACTA TGAAGAG) and OST 1600 (GCGCGAATTC TTAGAGCTTC GATTTCAGAC TTCCCTCGA) primers. The product was then reinserted into D3cpV using ClaI and EcoRI restriction sites. (pcDNA-)4mtD3cpV was a kind gift from A. Palmer and R. Tsien (Palmer et al., 2006) (Addgene #36324). D1cpV-px (PST 2169) was generated from the (pcDNA-) D1cpV (Palmer et al., 2004) (Addgene #37479) by amplifying an insert with oligonucleotide OST 2003 (GCGCGGATCC CATGGTGAGC AAGGGC) and OST 2002 (CGCGGAATTC TTAGAGCTTC GATTTCAGAC TTCCTATGAC AGGCTCGATG TTGTGGCGGA TCTTGAAGTT). The product was then reinserted into D1cpV using EcoRI and BamHI restriction sites. Pericam-px (PST 2170) was generated from ratiometric-pericam (for mitochondria) (Nagai et al., 2001) by amplifying an insert with OST 2116 (GCGCAAGCTT ATGAAGAGGCGC TGGAAGAAAA) and OST 2117b (GCGCGAATTC CTAGAGCTTC GATTTCAGAC TTCCTATGAC AGGCTTTGCT GTCATCATTT GTACAAACT), which was then re-inserted into ratiometric-pericam using EcoRI and HindIII restriction sites. (CMV-)R-GECO1 was a kind gift from R. Campbell (Zhao et al., 2011).

### Cells, cell culture and immunfluorescence

HeLa cells were cultured in low glucose Dulbecco’s Modified Eagle Medium (DMEM) medium (Biochrom) supplemented with 1% Pen/Strep (100units/ml Penicillin and 100 µg/ml Streptomycin), 1% (w/v) glutamine and 10% (v/v) Fetal Calf Serum (FCS) in 5% CO2 at 37°C. For immunofluorescent detection of PEX14, cells were fixed with 4% paraformaldehyde for 20 min, and permeabilized using 0.5% Triton X-100 in PBS for 5 min. After blocking for 30 min with 10% BSA in PBS (blocking buffer) at 37°C, antigens were labelled with primary antibodies at 37°C for 1 h. Rabbit anti-PEX14 (ProteinTech) primary antibody dilution in blocking buffer was 1:500. Labeling with the secondary antibodies conjugated to Cy3 (Life Technologies) was done for 1 h (1:500). Coverslips were mounted with ProLong Gold mounting medium with or without DAPI (Thermo Fisher Scientific).

NRCMs were isolated from newborn rats. Briefly, after the rats were sacrificed hearts were removed from the thoracic cavity, homogenized mechanically and digested in 1mg/ml collagenase type II containing calcium and magnesium-free PBS at 37°C with magnetic stirring. Supernatant was taken every 20 min and transferred to DMEM medium supplemented with Glutamax (Thermo Fisher Scientific), FCS and 1% Pen/Strep. Cells were then centrifuged, the cell pellet resuspended in fresh medium and transferred to a Petri dish for 45 min (37°C and 5% CO_2_). The fibroblasts adhered and NRCMs remained in the supernatant. NRCMs were then seeded on glass cover slips covered by Geltrex (Thermo Fisher Scientific).

Cells and cardiac differentiation of hiPSCs were described earlier (Borchert et al., 2017). Cells were studied 90 days after initiation of differentiation. Following differentiation, purity of hiPSC-CMs was determined by flow analysis (>90% cardiac TNT^+^) or by morphology (Borchert et al., 2017). HiPSC-CMs were maintained in RPMI 1640 supplemented with Glutamax, HEPES and B27 supplement.

### Ca^2+^ measurements

Cells (200,000 for HeLa and hiPSC-CMs and 500,000 for NRCMs) were seeded on glass cover slips and transfected with sensor plasmids using Effectene (Qiagen) (HeLa) or Lipofectamine LTX Reagent (Thermo Fisher Scientific) (hiPSC-CMs and NRCMs) according to the manufacturer’s instructions. Cells were imaged using a Zeiss Observer D1 (equipped with Zeiss Colibri 2 and Evolve 512 Delta EMCCD acquisition camera) or Axio Observer Z1 (equipped Zeiss Colibri 7, Definite Focus.2 and Zeiss Axiocam 702) with 40× oil Fluar (N.A. 1.3) objective at 37°C in a Ca^2+^-free imaging buffer (145 mM NaCl, 4 mM KCl, 10 mM HEPES, 10 mM glucose, 2 mM MgCl_2_, 1 mM EGTA, pH 7.4 at 37°C) 24 hours (HeLa and NRCMs) or 48 hours (hiPSC-CMs) after transfection. Where indicated, NRCMs were field-stimulated at 1 Hz with MyoPacer ES (IonOptix). Data were analyzed with AxioVision (Zeiss) and ZEN (Zeiss) software. Background and bleed-through (BT) were corrected in the FRET/donor ratio:

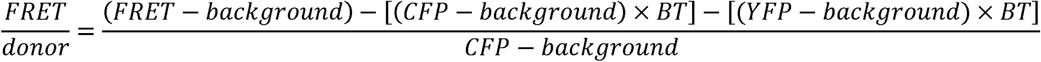

Excitation 420 nm and 505 nm with emission filters 483 ± 16 nm and 542 ± 14 nm, or excitation 438 ± 12 nm and 508 ± 11 nm with emission filters 479 ± 20 nm and 544 ± 14 nm were used. Where indicated, the concentration of Ca^2+^ in the imaging buffer was increased to 1 mM by doubling the buffer volume to the cells (e.g. during treatment with chemicals) by the addition of Ca^2+^-containing buffer (imaging buffer that contains 2 mM CaCl2 (pH 7.4, 37°C) instead of EGTA). HiPSC-CMs and NRCMs were incubated in 10 mM 2,3-butanedione monoxime (BDM) before the measurements. FRET ratios (calculated as FRET donor ratio) were calculated by subtracting the background intensity and correcting for crosstalk. ER-store depletion in the cells was induced by 100 μM histamine (HeLa) in Ca^2+^-free buffer or 1 μM Tg in Ca^2+^-containing buffer. For permeabilization, cells were treated with 0.01% digitonin in Ca^2+^-free EGTA buffer for 50 sec to 1 min and cytosol was washed out by rinsing twice with Ca^2+^-free EGTA buffer. Cell response to ionomycin was measured by the addition of 5 µM ionomycin in 10 mM Ca^2+^-containing buffer. Images for color LUT were made by applying Royal LUT on difference image of FRET and CFP in case of D3cpV-px and D1cpV-px, or difference image of 505 nm and 420 nm in case of pericam-px.

## Statistical analysis

Statistical significance was assessed using two-sided unpaired student’s t-test when comparing two groups, or one-way ANOVA followed by Tukey’s post hoc test when three groups were compared. Data were presented as Tukey’s box plots: the box is limited by 25^th^ and 75^th^ percentiles. Data points larger than 75^th^ percentile plus 1.5IQR (interquartile range) or smaller than 25^th^ percentile minus 1.5IQR are presented as outliers. All other data are covered by the whiskers.

## Author Contributions

ST conceived and designed the study. YS performed most experiments, conducted data analysis and prepared all figures. UB tested D3cpV-px localization, and measured Ca^2+^ in NRCMs and hiPSC-CMs. IB supervised Ca^2+^ measurements and provided access to the Zeiss Cell Observer Z1 with Colibri3 LED system. KSB provided hiPSC-CMs and contributed to manuscript writing. YS and ST wrote the manuscript. All authors read, revised and approved the manuscript.

## Acknowledgments

We thank Drs. Robert Campbell, Takeharu Nagai, and Nicolas Demaurex for providing GECO and pericam plasmids. We thank Julia Hofhuis for earlier work on peroxisomal calcium and for cloning of D3cpV-px and Xin Zhang for support with microscopy.

